# Regulating temperature and humidity inhibits the disease severity of tomato

**DOI:** 10.1101/2023.08.29.555345

**Authors:** Tianzhu Li, Jianming Li

**Author notes:** Corresponding author: Jianming Li E–mail address, College of Horticulture, Northwest Agricultural and Forestry University, Yangling 712100, China.

## Abstract

The environment significantly impacts the interactions between plants and pathogens, thus remarkably affecting crop disease occurrence. Regulating the environmental conditions in greenhouses to inhibit plant diseases is conducive to the development of green ecological agriculture. However, how to specific regulate the temperature and humidity to reduce plant diseases remain unclear. In this study, different methods of temperature and humidity regulation were performed. Histological observation and dual RNA–seq analysis showed that increasing temperature and decreasing humidity at 24 h post inoculation (hpi) for 12 h could effectively inhibit disease severity of tomatoes. The regulation induced the expression of heat shock proteins (HSPs) and defense–related genes of tomatoes in response to *B. cinerea*. Bcl–2–associated athanogenes (BAGs) may be involved in tomato resistance to *B. cinerea* mediated by temperature and humidity regulation. In addition, the infection process and the expression of toxin–related genes of *B. cinerea* such as sesquiterpene botrydial (BOT) and polyketide botcinic acid (BOA) was inhibited. Overall, we obtained the optimal regulation method of temperature and humidity to alleviate tomato gray mold, and preliminarily explored its mechanisms of inhibiting the disease.

## Introduction

Temperature and humidity play crucial roles in plant–pathogen interactions (Leisner et al., 2023; Martins et al., 2010; Saijo et al., 2019; Sarkar et al, 2011). Reducing plant diseases by regulating greenhouse temperature and humidity is becoming increasingly important in the field of ecological control (Hartmann et al. 2015; Padmini & Padmaja, 2010).

Temperature and humidity remarkably affect the infection process of pathogens and the defense response of plants. High air humidity is often associated with disease outbreaks (Miller et al., 2017; Yang et al., 2021), and plants are more susceptible to pathogens under high humidity (Miller et al., 2017). With increased air humidity, the virulence of *Sclerotinia* increases, and when humidity exceeds 80%, lettuce incidence is highest (Clarkson et al., 2014). The conidia of *B. cinerea* cannot germinate and the infection is inhibited when the relative humidity (RH) is lower than 80% (Blanco– Ulate et al., 2014). Increased temperature promotes the plant tissue necrosis, and facilitates the colonization of necrotrophic pathogens (Elad & Pertot, 2014). Changes in temperature considerably affect the growth, germination, survival, and virulence of pathogens (Kiewnick, 2006; Bugeme et al. 2008). Gray mold is a low–temperature, high–humidity disease. The optimum temperature for *B. cinerea* to cause diseases is 23°C, and high–humidity environments with relative humidity above 80% are conducive to disease occurrence (Eden et al., 1996). Previous studies have found that temperatures above 30°C can reduce the occurrence of gray mold (Benito et al., 1998; Amselem et al., 2011). Increasing the temperature of the tomato root region to 28°C can enhance the expression of defense genes and the eruption of reactive oxygen species (ROS), induce tomato immunity, and improve the resistance of tomato plants to *B. cinerea* (Gupta et al., 2021).

Tomato gray mold is a devastating plant disease caused by *B. cinerea*; it results in tomato rot and exerts 20%–40% damage, which can even reach 50%–70% during epidemic years (Cotoras & Silva, 2005). Temperature and humidity greatly affect the development of tomato gray mold. Our previous studies have shown that high temperature (35°C/28°C, day/night) and low humidity (RH 65%) can effectively alleviate the disease severity of tomatoes (Li et al., 2023). Therefore, in this study, we further explored how to specifically regulate temperature and humidity to inhibit plant diseases. We hope to achieve disease control through reasonable regulation of greenhouse environmental conditions, and finally realize the development of green ecological agriculture.

## Results

### The regulation of temperature and humidity inhibits the disease severity of tomato plants

First of all, we conducted an experiment on when to start regulating temperature and humidity, and we found that regulating temperature and humidity at 24 h post inoculation (hpi) could effectively inhibit tomato gray mold (Supplemental Figure S2). Further, we took 24 hpi as the premise to explore how long the regulation would be the best for inhibiting the disease. Among all the treatments, the disease of tomato plants in the 10 h treatment was the most serious, and leaf lesions expanded rapidly with the increase of time (Figure 2A). The infected tomato leaves in the control treatment became yellow and brown at 3 cycles (Figure 2A, c5). We found that compared with the control, the 6 h, 8 h, and 12 h treatments could effectively inhibit the spread of disease lesions and disease severity of tomatoes (Figure 2, B, C, and E), and the 12 h treatment reduced gray mold severity the most (Figure 2, A and E). The disease inhibition effect of tomato plants under different treatments followed the order: 12 h > 8 h > 6 h > control > 10 h.

**Figure 1.**
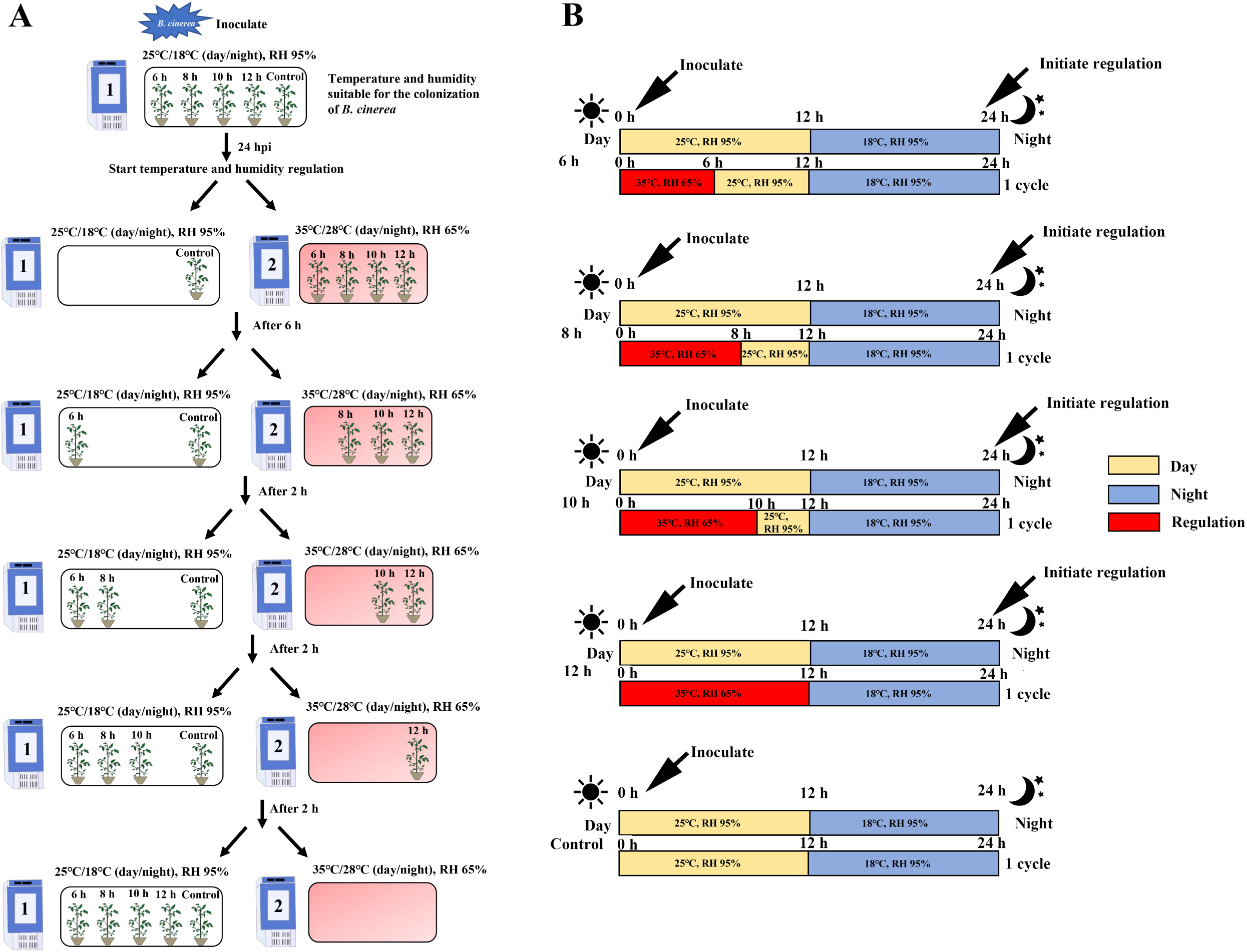
Schematic diagram of temperature and humidity regulation. A: Illustration of tomato plants moving between two incubators under different treatments; B: Environmental conditions of tomato plants in the 6 h, 8 h, 10 h, 12 h, and control treatments. The figure shows one cycle.

**Figure 2.**
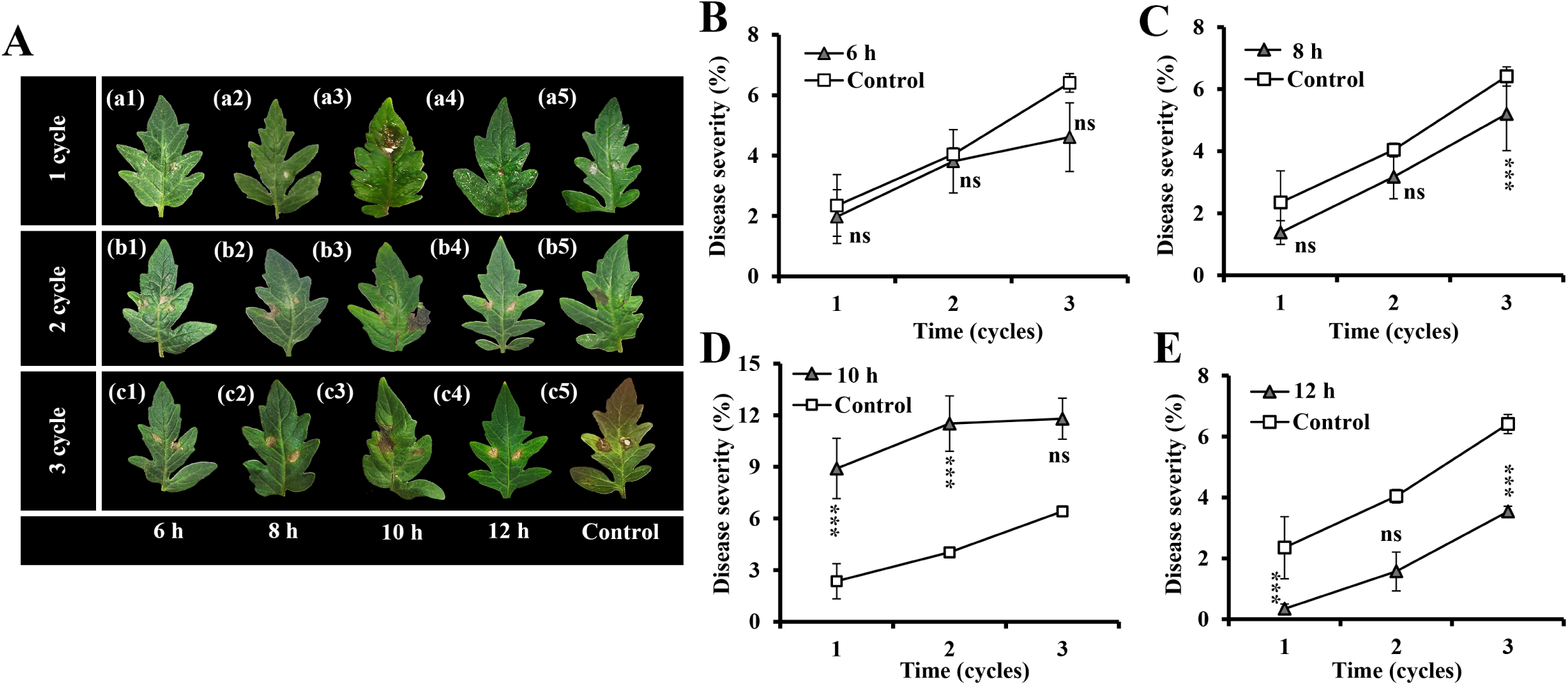
The disease severity of tomatoes in the different treatments at 1–3 cycles. A: Characterization of infected tomato leaves under different treatments at 1–3 cycles; B, C, D, E: Disease severity of tomato plants in the 6 h, 8 h, 10 h, and 12 h treatments at 1–3 cycles. Graphs represent the results of at least three repetitions ± SEM. Results were analyzed for statistical significance by using multiple t–tests (one per time point). Asterisks represent statistically significant differences. ***P < 0.01; **P < 0.05; ns: not significant.

### The regulation of temperature and humidity inhibits the infection process of *B. cinerea*

Among the five treatments, *B. cinerea* grew best with the largest colony diameter in the control (Figure 3A), and a large number of mycelia were observed in tomato leaves at 1 cycle (Figure 3C). The colony diameter of *B. cinerea* treated for 6 h was significantly larger than that treated for 8 h, 10 h and 12 h (Figure 3B), whereas the growth of mycelia in leaves was less (Figure 3C, a2, b2, and c2). The colony diameter of the 10 h treatment was smaller, whereas a large number of mycelia were observed in the tomato leaves at 1 cycle (Figure 3C, a4). Among the five treatments, 12 h inhibited the *B. cinerea* growth and infection process in tomatoes the most.

**Figure 3.**
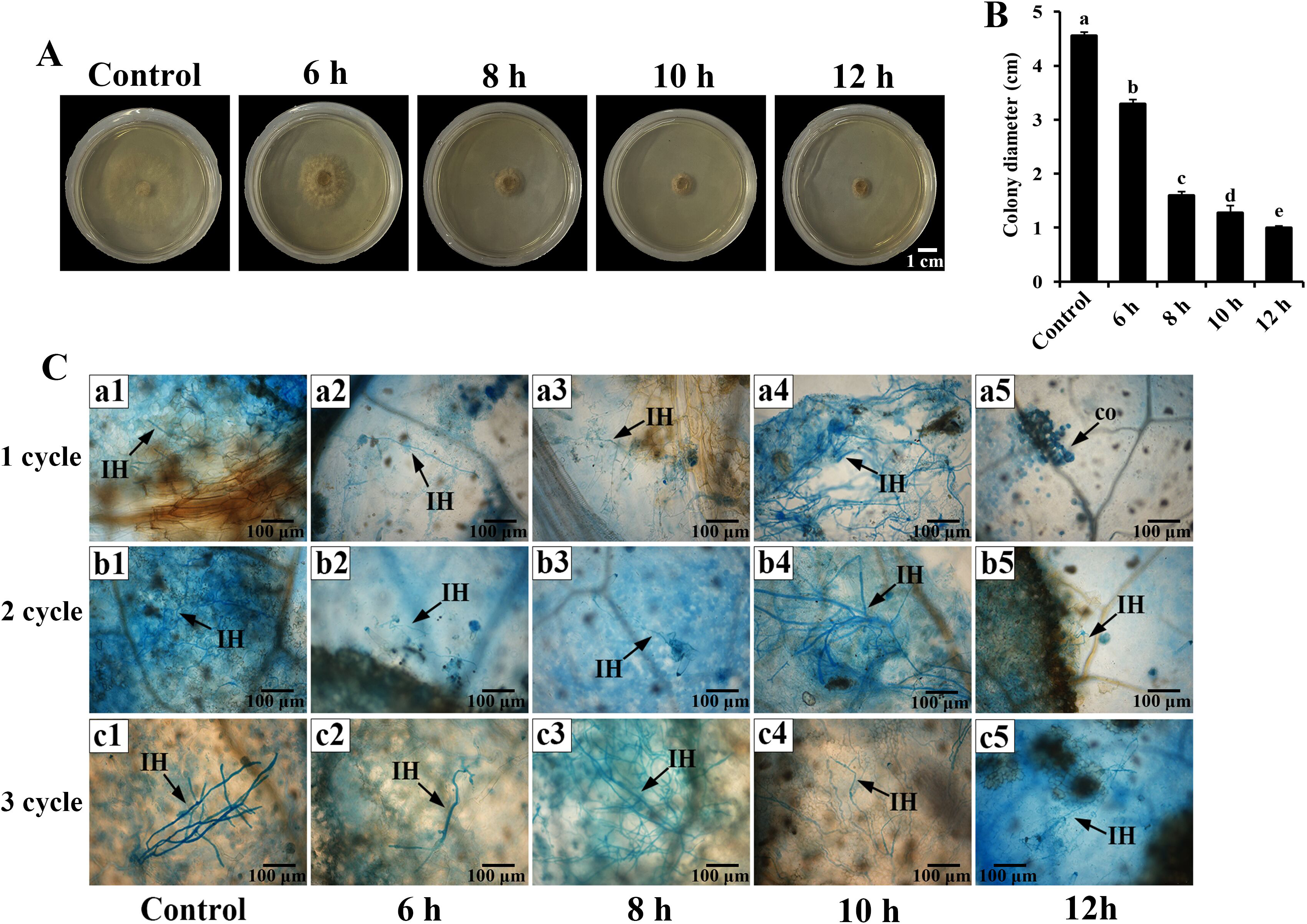
Growth and development of *B. cinerea* in the different treatments. A: Effects of different treatments on the growth of *B. cinerea* in vitro at 3 cycles; B: Colony diameter of *B. cinerea* growing in vitro under different treatments at 3 cycles; C: The infection process of *B. cinerea* in tomato leaves under different treatments at 1–3 cycles. Graphs represent the results of at least three repetitions ± SEM. Results were analyzed for statistical significance by using multiple t–tests. Lowercase letters on the error bar represent significant difference at the 0.05 level of different treatments. co: conidium; IH: infection hyphae. Scales = 100 μm.

### The regulation of temperature and humidity induces the expression of heat shock protein genes of infected tomatoes

We analyzed the dual–RNA seq data of the 12 h (the treatment that most effectively inhibits the disease) and control (the treatment that most suitable for the development of the disease) treatments to explore the effect of regulating temperature and humidity on the interactions between tomato and *B. cinerea*. A total of 4242 Differentially expressed genes (DEGs) of infected tomato plants were detected in 12 h vs. control, of which 2368 genes were up–regulated and 1874 genes were down– regulated (Supplemental Table S2 and S3). Mapman visualized the individual genes in plant biotic stress pathway (Supplemental Figure S3). The abiotic stress, heat shock proteins (HSPs) and defense genes were greatly upregulated (Supplemental Figure S3). Thirty–nine genes with log FC >2 related to the HSPs were selected for analysis (Figure 4, A and B; Supplemental Table S4). *HSP20*, *HSP70* and *HSP90* were mainly involved in the response of tomato to *B. cinerea* under regulating in temperature and humidity (Figure 4B). To further verify whether the upregulation of *HSPs* of tomatoes was caused by environment or pathogens, we measured the gene expressions of *HSP17.4*, *HSP20*, *HSP70*, and *HSP90* in inoculated and uninoculated tomato plants in 12 h and control treatments (Figure 4, C–F). Temperature and humidity regulation induced the expression of *HSP17.4*, *HSP70*, and *HSP90* in uninoculated tomatoes. *B. cinerea* infection did not induce the expression of *HSP70* and *HSP90* in tomatoes either under control or 12 h treatments. However, regulation significantly up– regulated *HSP17.4*, *HSP20*, and *HSP90* in inoculated tomato plants, and the expression of *HSP20* in inoculated tomatoes was higher than that in uninoculated tomatoes. In general, we found that pathogen infection almost did not induce the expression of *HSPs* in tomatoes, and temperature and humidity regulation induced the upregulation of *HSP17.4*, *HSP20*, and *HSP90* in infected tomato plants. The upregulation of *HSP20* was mainly caused by regulating in temperature and humidity.

**Figure 4.**
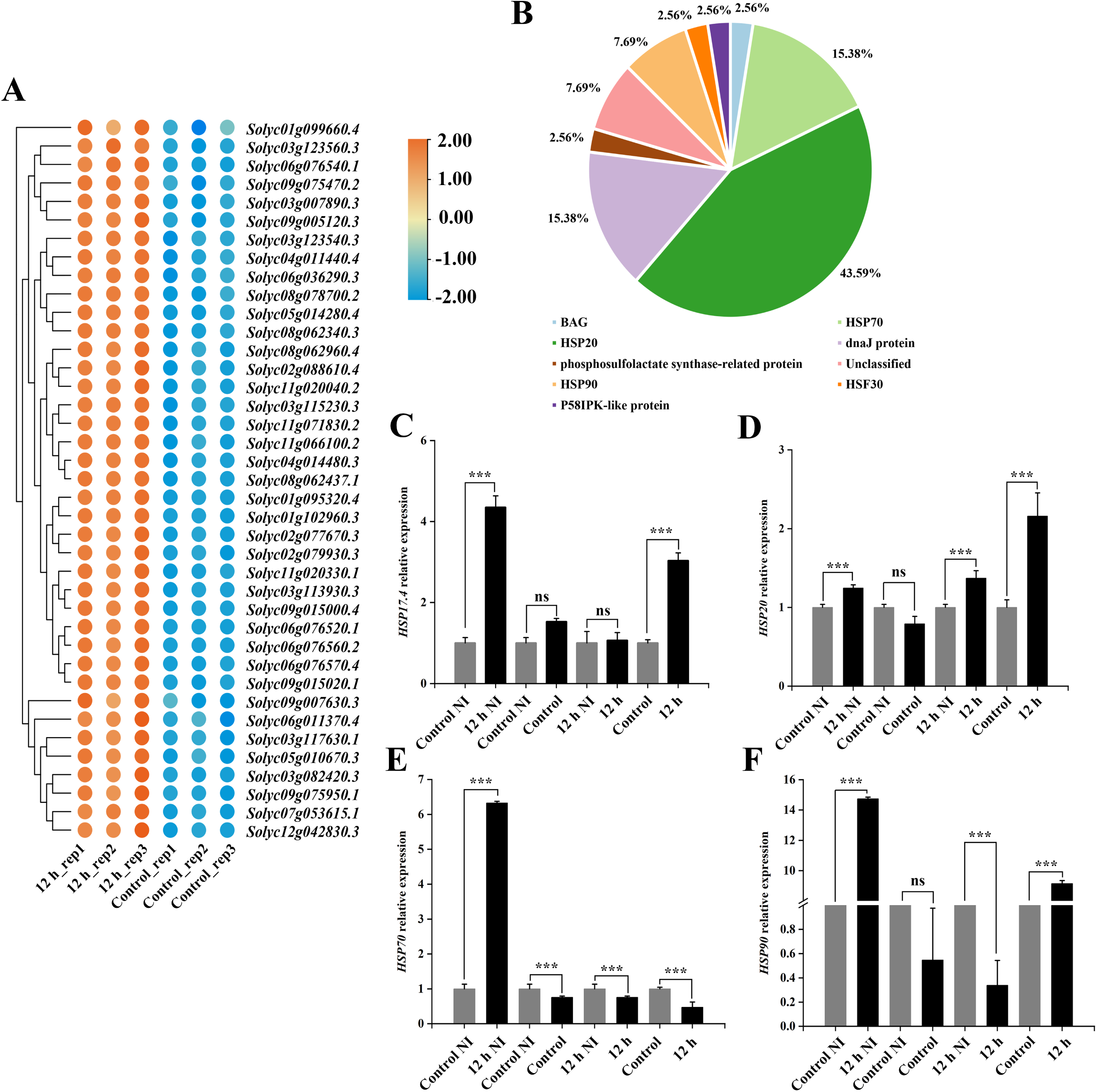
Changes of heat shock proteins (HSPs) related genes of tomato plants infected by *B. cinerea* in 12 h vs. control at 1 cycle. A: Heat map of HSPs related genes (log FC >2) expression in infected tomato plants. The relative level of abundance is indicated by a color gradient from low (blue) to high (orange). B: Type and proportion of upregulated HSPs genes in 12 h vs. control. C, D, E, F: The expression of *HSP17.4, HSP20, HSP70*, and *HSP90* genes in inoculated and uninoculated tomato plants in the 12 h and control treatments. Graphs represent the results of at least three repetitions ± SEM. Results were analyzed for statistical significance by using multiple t–tests under different treatments. Asterisks represent statistically significant differences. ***P < 0.01; **P < 0.05; ns: not significant.

### The regulation of temperature and humidity induces the expression of defense genes of infected tomatoes

In 12 h vs. control, thirty–four tomato DEGs related to the pattern recognition receptors (PRRs) were detected, of which nine genes were down–regulated and twenty–five genes were up–regulated (Figure 5A and Supplemental Table S5). The 12 h treatment induced the expression of *LRR*–*RLK EFR*, *RLK*–*HSL1*, *LRR*–*RLK FEI2*, *LRR*–*RLK HSL2*, *LRR*–*RLK ERECTA,* and *LRR*–*RLK FLS2* of infected tomatoes (Figure 5, A and B). Eleven Ca^2+^ sensors genes with EF–hand structure were up– regulated in infected tomatoes in 12 h vs. control (Figure 5C and Supplemental Table S6). Subsequently, we measured calmodulin (CaM) content of inoculated and uninoculated tomato plants in the 12 h and control treatments during 1–3 cycles (Figure 5D). The results showed that temperature and humidity regulation increased CaM content in uninoculated plants at the early stage, then decreased. Regulation induced an increase of CaM content in infected tomato plants at 3 cycle.

**Figure 5.**
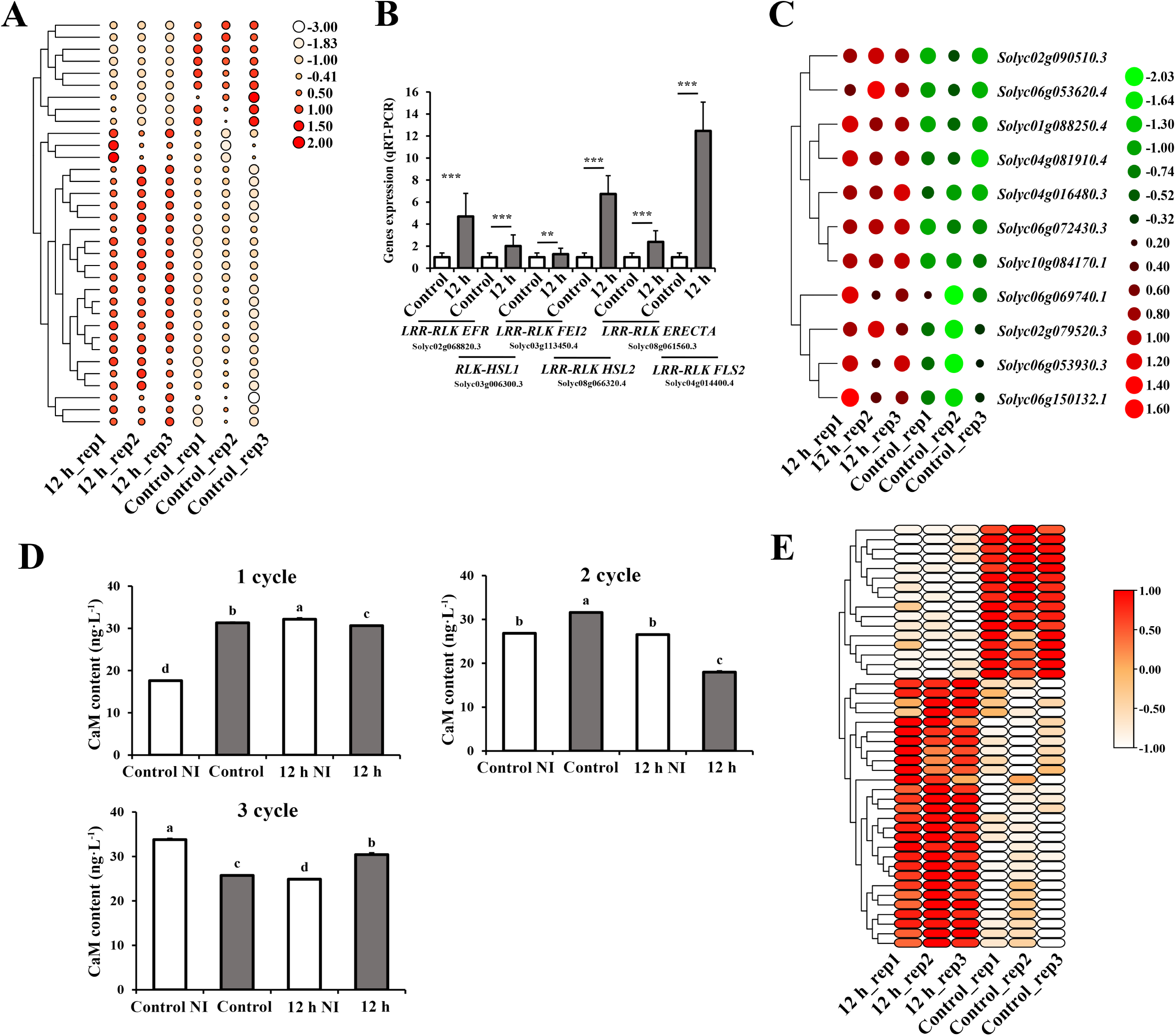
Changes of defense related genes of tomato plants infected by *B. cinerea* in 12 h vs. control. A: Heat map of pattern recognition receptors (PRRs) genes expression in infected tomato plants in 12 h vs. control. The relative level of abundance is indicated by a color gradient from low (white) to high (red). B: Genes expression (qRT–PCR) of some *PRRs* are up–regulated in infected tomato plants in the 12 h treatment; C: Heat map of Ca^2+^ sensors genes with EF–hand structure expression in infected tomato plants in 12 h vs. control. The relative level of abundance is indicated by a color gradient from low (green) to high (red). D: Calmodulin (CaM) contents of inoculated (12 h, control) and uninoculated (12 h NI, control NI) tomato plants at 1–3 cycles. E: Heat map of defense genes expression in infected tomato plants in 12 h vs. control. The relative level of abundance is indicated by a color gradient from low (white) to high (red). Graphs represent the results of at least three repetitions ± SEM. Results were analyzed for statistical significance by using multiple t–tests under different treatments. Asterisks represent statistically significant differences. ***P < 0.01; **P < 0.05; ns: not significant. Lowercase letters on the error bar represent significant difference at the 0.05 level of different treatments.

Forty–four tomato defense genes were differentially expressed in 12 h vs. control, of which twenty–seven genes were up–regulated and seventeen genes were down– regulated (Figure 5E and Supplemental Table S7). Further analysis of defense genes showed that all five genes encoding pathogenesis–related (PR) proteins were down– regulated, and most of the up–regulated defense genes encoding disease resistance proteins, receptor–like kinase (RLK) and receptor–like proteins (RLPs) (Supplemental Table S6).

### *BAGs* are involved in the resistance of tomato to *B. cinerea* under regulating in temperature and humidity

Analysis of *HSPs* and Ca^2+^ sensors genes of inoculated tomato plants in 12 h vs. control revealed that both genes contained BAG protein genes (Figure 4 and 5; Supplemental Table S4 and S6). Therefore, we further analyzed the tomato DEGs encoding BAG proteins in 12 h vs. control. Five genes encoding BAG protein in 12 h vs. control (Figure 6A and Supplemental Table S8). Among them, four up–regulated *BAG* genes each contain a BAG domain, and two genes also contain an IQ calmodulin binding motif. To further study whether the up–regulation of *BAGs* in tomatoes was caused by environment or pathogens, we measured all *BAGs* family members (*BAG 1∼1*0) in tomatoes (Figure 6, B–K). Regulating temperature and humidity induced the expression of *BAG 1, BAG 6, BAG 7, BAG 8* and *BAG 9* in uninoculated tomato plants. The expression of *BAG 1, BAG 3* and *BAG 9* was induced under control treatment, and the expression of *BAG 3* and *BAG 10* was induced under regulating temperature and humidity conditions when pathogen infection. Regulation induced the expression of *BAG 6∼9* in inoculated tomatoes. It is noteworthy that *BAG 6, BAG 7* and *BAG 8* were all involved in plant responses to environment and environment– plant–pathogen interactions, and the expression levels of these genes in the latter (response to environment–plant–pathogen interactions) were higher than those in the former (response to environment).

**Figure 6.**
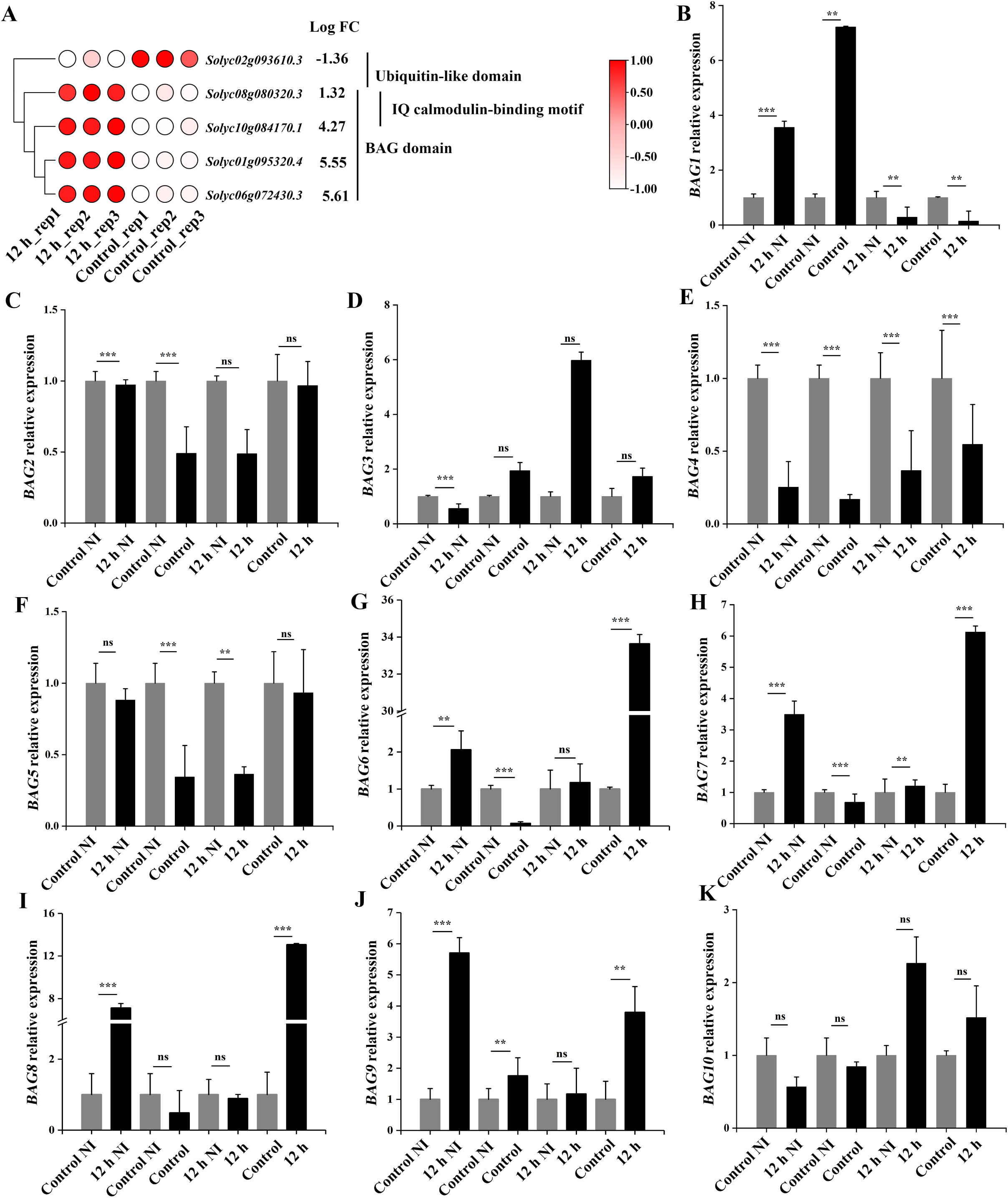
Bcl–2–associated athanogenes (BAGs) expression level in tomato plants in the 12 h and control treatments. A: Heat map of *BAGs* expression in infected tomato plants in 12 h vs. control. The relative level of abundance is indicated by a color gradient from low (white) to high (red); B–K: *BAG1∼10* expression levels of inoculated and uninoculated tomato plants in the 12 h and control treatments at 1 cycle. Graphs represent the results of at least three repetitions ± SEM. Results were analyzed for statistical significance by using multiple t–tests under different treatments. Asterisks represent statistically significant differences. ***P < 0.01; **P < 0.05; ns: not significant.

### The regulation of temperature and humidity inhibits the expression of genes related to toxin synthesis of *B. cinerea*

Dual RNA–seq analysis of inoculated tomato plants in 12 h vs. control showed that a total of 1623 DEGs of *B. cinerea* were detected, of which 917 were up–regulated and 706 were down–regulated (Supplemental Table S9 and S10). Among the 706 down–regulated DEGs of *B. cinerea*, 32 GO terms were significantly enriched, and most enrichment enriched GO term was related to the pathogenesis of *B. cinerea* (Supplemental Figure S4A and Table S11). Among the 11 down–regulated pathogenesis genes of *B. cinerea*, nine genes encoding botcinic acid (BOA) and two genes encoding botrydial (BOT) (Supplemental Figure S4B and Table S11).

### Gene expression detection by quantitative real–time–PCR

To verify the results of the dual RNA–seq assay, quantitative real–time–PCR (qRT–PCR) was conducted to analyze the relative expression of six tomato genes and nine *B. cinerea* genes. The results showed that the expression patterns of these representative genes were consistent with the transcriptome data (Supplemental Figure S5). This result indicates that transcriptome data are reliable.

## Discussion

Temperature and humidity play indispensable roles in plant–pathogen interactions (Martins et al., 2010; Sarkar et al, 2011). Reducing disease severity by regulating temperature and humidity is of great significance to “green agriculture” (Padmini & Padmaja, 2010; Hartmann et al., 2015). Based on our previous study, it was found that high temperature (35°C/28°C, day/night) and low humidity (RH 65%) effectively inhibit tomato gray mold (Li et al., 2023). In this study, we further solved the problems of how specific implementation of high temperature and low humidity regulation can effectively alleviate gray mold of tomatoes. Regulation at 24 hpi for 12 h effectively inhibit the disease severity of tomatoes and the infection ability of *B. cinerea*. The regulation induced the expression of *HSPs*, *PRRs*, *BAGs*, and defense genes of tomato plants, inhibited the expression of BOT and BOA genes of *B. cinerea*. The tomato *BAG6∼8* may be involved in the interactions between plants and *B. cinerea* under the regulation of temperature and humidity.

Histological and microscopic observations showed that all treatments except 10 h could inhibit the disease severity of tomato plants, and the 12 h treatment had the best inhibitory effect (Figure 2), while the growth and infection process of *B. cinerea* were inhibited in the 10 h treatment (Figure 3), indicating that the increase of tomato gray mold caused by the 10 h treatment may be due to the decline of plant resistance caused by environmental stress. Further, we performed dual–RNA seq analysis between the 12 h and control treatments, we found that tomato *HSPs* were greatly induced in 12 h vs. control (Figure 4). HSPs can be induced by high temperature and other environmental cues (cold, drought, salinity, etc.) (Yu et al., 2016), and many HSPs respond not only to abiotic stress, but also to pathogen infection (Bhattarai et al., 2007; Breiman, 2014). In recent years, HSPs in plants have attracted extensive attention due to their new functions in innate immunity (Arofatullah et al., 2019; Kharisma et al., 2022). In our study, the 12 h treatment induced up–regulation of *HSP20*, *HSP90*, and *HSP17.4* in the infected plants, and the expression of *HSP20* was mainly caused by environment–plant–pathogen interactions (Figure 4). HSP90 is the most abundant cytoplasmic HSP family in eukaryotic and prokaryotic cells, which can rapidly induce responses to various stress conditions (Breiman, 2014). As a positive regulator of plant immunity, HSP90 can directly interact with R protein, and its substrates are mostly kinases and transcription factors that activate defense responses (Sangster & Queitsch, 2005). We speculate that *HSP90* might be involved in the defense response of tomato to *B. cinerea* mediated by regulating in temperature and humidity. HSP20 is a representative small heat shock protein (sHSP), which is the most important and abundant protein in many species under thermal stimulation (Yu et al., 2016). *HSP20* in tabacum is involved in the plant disease resistance (Maimbo et al., 2007), and *SlHSP20* has a potential role in mediating tomato response to environmental stress (Yu et al., 2016). The 12 h treatment induced the expression of *HSP20*, and the expression level of *HSP20* in inoculated plants was higher than that in uninoculated plants, suggesting that the upregulation of *HSP20* may be caused by environmental changes and pathogen infection, and participated in the process of temperature and humidity regulation of tomato gray mold.

Many tomato genes encoding PRRs were upregulated in 12 h vs. control (Figure 5). PRRs of host plants can recognize the pathogen–associated molecular patterns (PAMP) released by pathogens and cause a series of physiological and biochemical changes in plant cells to resist the infection of pathogens (Zipfel, 2014; Bohm, 2014). The increased *PRRs* expression is associated with enhanced immune output, with cell damage, sufficient to produce a strong local response to invading pathogens (Zhou et al., 2020). Recognition of PRRs and PAMP results in changes in ion fluxes and hormone levels, which in turn induce transcription of plant defense genes (Hann & Rathjen, 2007; Robatzek et al., 2007; Takai et al., 2008). In 12 h vs. control, many Ca^2+^ sensor genes with EF–hand structures of tomatoes were induced (Figure 5).

CaMs are an important member of the Ca^2+^ sensor family (Perochon, et al., 2011; Reddy, et al., 2011; Boudsocq et al., 2013). The environment increases CaM contents in uninoculated plants at the early stage, then decreased; the pathogens induce the increase and then decrease of CaM content in tomatoes under control conditions, and first decrease and then increase under regulating temperature and humidity conditions (Figure 5D). These results indicate that the abiotic stress caused by temperature and humidity regulation in a short period of time led to Ca^2+^ influx in plants and increased intracellular Ca^2+^ content. In addition, we found that many tomato defense genes were upregulated in 12 h vs. control, including genes encoding RLK, RLPs, and disease resistance proteins (Figure 5). Notably, all the DEGs encoding PR proteins in tomatoes were downregulated. Expression of *PR* genes is often involved in SA– mediated plant systemic acquired resistance (SAR) (Pucciariello et al., 2017). However, our study found that the temperature and humidity regulation mediated immune response of tomato to *B. cinerea* did not induce the expression of *PR* genes. Earlier studies found that the effects of high temperature in the greenhouse involved an indirect systemic effect that made the host less susceptible to disease, and that the effect was also observed in harvested branches that were no longer heat–treated, and that the effect was systemic (Elad et al., 2014). We speculate that in the tomato–*B. cinerea* pathological system, the enhancement of tomato’s defense ability against *B. cinerea* by temperature and humidity regulation maybe belongs to induced resistance (IR), and this IR is systematic.

In 12 h vs. control, tomato *BAGs* were identified in both HSPs and Ca^2+^ sensors genes (Figure 4 and 5), and four *BAGs* were up–regulated (Figure 6). The BAG family is a highly conserved family of helper molecular chaperone proteins, and BAG proteins in plants play an important role in a series of apoptosis processes such as pathogen invasion, abiotic stress and plant development (Takayama & Reed, 2001). Currently, seven and ten BAG family members have been identified in *Arabidopsis* and tomato, respectively. The calmodulin–binding domain contained in *AtBAG5∼7* in *Arabidopsis* is plant–specific, suggesting that BAGs may be involved in an important process of calcium signal transduction (Thanthrige et al., 2020). In our study, *BAG6∼8* were significantly induced in the inoculated tomato plants after temperature and humidity regulation. This result suggests that *BAG6∼8* may be involved in the temperature and humidity regulation–mediated defense response of tomato to *B. cinerea*. Early studies showed that *BAG8* and *BAG9* in tomatoes have a positive effect on heat stress, and they interacted with *CaM1∼6* and *HSP20* (Shemetov & Gusev, 2011). *AtBAG6* may be involved in the basic resistance of *Arabidopsis thaliana* to necrotrophic pathogens such as *B. cinerea* (Li et al., 2016). We speculate that the increased resistance of tomato to *B. cinerea* mediated by regulating in temperature and humidity may be related to the increased expression of *BAG6*. AtBAG7 is one of the most characteristic BAGs, is an endoplasmic reticulum (ER) localization protein that plays a central regulatory role in heat–induced UPR pathway (Li et al., 2017). AtBAG7 interacts with *WRKY29* to promote plant resistance (Li et al., 2017). However, we did not find that *WRKY29* was induced in 12 h vs. control, which may indicate that *BAG7* is involved in plant resistance through other pathways. In general, we speculate that *BAG6∼8* have a positive regulatory role in the temperature and humidity regulation–mediated defense response of tomato against *B. cinerea*, and might be involved in the expression of *HSPs* and Ca^2+^ signal transduction of tomatoes.

Many *B. cinerea* genes encoding BOT and BOA were down–regulated in 12 h vs. control (Supplemental Figure S4). BOT and BOA are two main plant toxins produced by *B. cinerea* (Dalmais et al., 2011). We found that *Bcbot1* and *Bcbot3∼5* in *B. cinerea* were down–regulated by regulating in temperature and humidity. Previous studies showed that deletion of *Bcbot1* encoding monooxygenase reduced virulence of *B. cinerea* (Dalmais et al., 2011). Temperature and humidity regulation may reduce the infection ability of *B. cinerea* by inhibiting pathogen virulence. In addition, fungal genes encoding *Bcboa2*, *Bcboa6∼8, Bcboa11*, and *Bcboa13* were down–regulated. The BOA is a phytotoxic polyketone involved in the virulence of *B. cinerea*, and pathogens use toxic secondary metabolites as molecular weapons to induce plant cell death (Porquier et al., 2019). *Bcboa13* is an important nucleoprotein regulating the biosynthesis of BOA, and plays a positive role in the expression of other *Bcboa1∼12* genes (Porquier et al., 2019). We speculate that the down–regulation of toxin–related genes such as *BOT* and *BOA* produced by *B. cinerea* might be related to the regulation of temperature and humidity, which reduced the virulence of *B. cinerea* and inhibited the ability of pathogen infection. Previous studies found that the production of fungal secondary metabolites is strictly controlled by environmental signals, and deviation from most suitable environmental conditions for pathogen growth and development may lead to the inhibition of the expression of virulence genes (Brakhage, 2012). This result is consistent with our research conclusions.

## Conclusion

In conclusion, we obtained that temperature and humidity regulation at 24 hpi for 12 h could effectively inhibit the disease severity of tomatoes and infection ability of *B. cinerea*. The regulation induced the expressions of *HSPs*, *PRRs*, *BAGs*, and defense genes in infected tomato plants, and inhibited the expressions of genes related to the pathogenesis of *B. cinerea*. The tomato *BAG6∼8* may participate in the resistance of plants to *B. cinerea*.

## Materials and methods

### Material and treatment

Experiments were conducted at the Laboratory of Physiology, College of Horticulture of Northwest A&F University, China. The tomato variety Jinpeng No. 1 was purchased from Yangling Yufeng Seed Industry, Shaanxi, China. Tomatoes were cultured at 25°C/18°C (day/night), RH of 65%, and photoperiod of 12 h/12 h (light/dark) for seven weeks. *B. cinerea* was isolated from diseased tomato fruits and grown on potato dextrose agar (PDA) at 26°C for seven days prior to inoculation. The conidia suspension concentration was adjusted to 1×10^6^ conidia per mL by using a blood–cell counting plate. Tomato leaves were inoculated with 20 μL liquid drops, with 2 drops on each leaf (Windram et al., 2012).

Temperature and humidity regulation mean that the temperature is increased to 35°C/28°C (day/night) and the humidity is reduced to 65%. The aim is to obtain the optimal method of temperature and humidity regulation, so we have designed the experiment of when to regulate and how long to regulate. Regulating at 12 hpi, 18 hpi and 24 hpi respectively to obtain the optimal regulation time point (Supplemental Figure S1). Then, on this basis, the regulation was performed for 6 h, 8 h, 10 h and 12 h respectively (Figure 1). All experiments were carried out in incubators with precise temperature and humidity regulation. In order to clearly observe the inhibitory effect of regulation on the disease, we set the unregulated treatment as the control, that is, the temperature of 25°C/18°C (day/night) and the humidity of 95%, which are suitable conditions for the occurrence of gray mold. Each treatment placed 20 tomato plants, 10 of which were inoculated and 10 of which were not inoculated, to rule out the effects of environmental changes on the plants. From the start of regulating to the subsequent 24 h as a cycle, a total of 3 cycles were executed.

### Gray mold disease evaluation

Disease severity was calculated as the percentage of the leaf lesion area over the entire leaf area (Meller Harel et al., 2014). The diseased leaves were placed on a white paper with scale line, photographed, and measured using Image J software. The area under the disease progress curve (AUDPC) was used to assess the susceptibility of the plant and expressed as the percentage of infection over a period of time. The higher AUDPC is, the higher the susceptibility of the plant is to the disease (Bonierbale et al., 2007). The experiment was conducted independently three times. AUDPC is usually calculated using the formula.

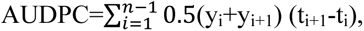

where t_i_ is the date of investigation, y_i_ is the percentage of infected leaves during each investigation, and n is the total number of days.

### Observation of the infection process of *B. cinerea*

The growth of *B. cinerea* under different treatments was observed to study the effect of environment on pathogens. *B. cinerea* was inoculated on PDA, and after three cycles, colony diameters of *B. cinerea* under different treatments were measured and photographed. The infection process of *B. cinerea* in tomato leaves under different treatments was observed by using a light microscope. The leaves at the third leaf position were selected from each treatment and stained with lactate phenolic cotton orchid dyeing solution. The experiment was performed independently three times (Cheng et al., 2012).

### Determination of calmodulin content in tomato plants

The contents of CaM in tomato leaves were determined by the Zoonbio Biotechnology Co. Ltd, Nanjing, China.

### RNA extraction and data preprocessing

Total RNAs were extracted from the 12 h and control treated tomato leaves after one cycle by using the TRIzol Reagent (Invitrogen, cat. no 15596026). DNA digestion was carried out after RNA extraction by DNaseI. RNA quality was determined by examining A260/A280 with a NanoDrop^TM^ OneC spectrophotometer (Thermo Fisher Scientific Inc). RNA integrity was confirmed via 1.5% agarose gel electrophoresis. The qualified RNAs were quantified using Qubit3.0 from the Qubit^TM^ RNA Broad range Assay Kit (Life Technologies, Q10210). Transcriptome sequencing was completed on the Illumina sequencing platform. An Illumina PE library (∼300bp) was constructed and sequenced. The quality of the obtained sequencing data was controlled, and the transcriptome data were analyzed by bioinformatics methods. Raw data were cleaned using Trimmomatic software (version 0.36), and the clean data were compared with tomato (https://solgenomics.net/) and *B. cinerea* (http://fungi.ensembl. org/Botrytis_ cinerea/Info/Index) genomes to obtain comprehensive transcript information.

### RNA–Seq data analysis

The reads per kilobase per million reads (RPKM) value was used as the measurement index to obtain gene expression statistics. The absolute value of log FC > 1 was adopted, and *P* value< 0.05 was used as the standard for gene differential expression to indicate that the gene was a differential expression gene (DEG). Subsequently, cluster analysis was conducted for DEGs, and different expression regulation patterns were classified via hierarchical clustering. The Gene Ontology database (http://www.geneontology.org) was utilized to classify one or a set of genes involved in accordance with the genes’ biological process (BP), molecular function (MF), and cellular component (CC). Metabolic pathway analysis of DEGs was performed using the Kyoto Encyclopedia of Genes and Genomes (KEGG; http://www.genome.jp/kegg/).

### qRT–PCR analysis

2 μg RNA was converted to first–strand cDNA by using reverse transcriptase (Promega, Madison, WI) and Oligo–D (T) primers in accordance with the manufacturer’s instructions. qRT–PCR was performed under the Power SYBR Green Master Mix protocol (Life Technologies, Rockville, MD) by using a rotor–gene Q machine (Qiagen, Valencia, CA). The specific primers used in this study are shown in Supplemental Table S1. Relative expression quantification of gene expression was performed using the △△CT method (Fernandez et al., 2021).

### Data analysis

All experimental data were presented as average ± SEM. Differences among different groups were analyzed for statistical significance with a one–way ANOVA in SPSS software (ver. 21.0) (Gabler et al., 2003; Poolsawat et al., 2012). Differential gene expression was inferred based on the total mapping counts using the Edge R package (version 2.6.8) implemented in R. GO and KEGG enrichment analyses were implemented for the DEGs by KOBAS software (version 2.1.1) with a *P*–value cutoff of 0.05 to judge statistically significant enrichment. MapMan was used to visualize individual genes involved in the global metabolic pathways. Moreover, a heat map was constructed with the software TBtools (version 0.53). Figures were created in Microsoft Excel (ver. 2019, Microsoft Corporation, Redmond, WA). All images were combined in Adobe Photoshop (CC 2018).

## Supplementary data

**Supplemental Figure S1.** Schematic diagram of 12 hpi, 18 hpi, and 24 hpi treatments. A, One cycle of the12 hpi treatment. Regulation began (start the experiment of changing temperature and humidity) at the 12 hpi and lasted for 12 h, the plants were then moved back to the previous condition (25℃/18℃ (day/night) with RH 95%) for 12 hours. 24 h was a cycle, and the experiment was conducted for three cycles. B, One cycle of control (25℃/18℃ (day/night) with RH 95%, the treatment was not be regulated). C, One cycle of the 18 hpi treatment. D, One cycle of control. E, One cycle of the 24 hpi treatment. F, One cycle of control. Each treatment had an unregulated control group. Each treatment was repeated three times.

**Supplemental Figure S2.** Disease severity of tomato plants in the 12 hpi, 18 hpi, and 24 hpi treatments at 1–3 cycles. A and B, Disease severity and area under the disease progress curve (AUDPC) in tomato plants in the 12 hpi treatment. C and D, Disease severity and AUDPC in tomato plants in the 18 hpi treatment. E and F, Disease severity and AUDPC in tomato plants in the 24 hpi treatment. Graphs represent the results of at least three independent experiments ± SEM. Results were analyzed for statistical significance using multiple t–tests (one per time point). \*\*\**P*<0.01; \*\**P*<0.05; ns: non–significant. Asterisks represent statistically significant differences.

**Supplemental Figure S3.** Overview of biotic stress of tomato plants in 12 h vs. control visualized by using MapMan. The genes that were significantly up regulated–and down–regulated are shown in red and blue, respectively. The scale bars display Log 2 (fold changes).

**Supplemental Figure S4.** Genes related to pathogenesis of *B. cinerea* are downregulated in 12 h vs. control. A, Enriched GO terms of downregulated *B. cinerea* genes. B, Heat map of pathogenesis–related genes expression of *B. cinerea* in 12 h vs. control. The relative level of abundance is indicated by a color gradient from low (green) to high (red).

**Supplemental Figure S5.** qRT–PCR of genes in tomato and *B. cinerea*. The bar graph represents the gene expression from RNA–seq, and the line graph represents the gene expression from qRT–PCR. Three biological replicates were used to calculate the mean and the standard error of the mean.

**Supplemental Table S1** qRT–PCR primers used in this study.

**Supplemental Table S2** Upregulated DEGs of tomatoes in 12 h vs. control.

**Supplemental Table S3** Downregulated DEGs of tomatoes in 12 h vs. control.

**Supplemental Table S4** DEGs with log FC >2 of tomatoes in HSPs in 12 h vs. control.

**Supplemental Table S5** DEGs of tomatoes in PRRs in 12 h vs. control.

**Supplemental Table S6** DEGs of tomatoes in Ca^2+^ sensors with EF–hand structure in 12 h vs. control.

**Supplemental Table S7** DEGs of tomatoes in defense genes in 12 h vs. control.

**Supplemental Table S8** DEGs of tomatoes in BAGs in 12 h vs. control.

**Supplemental Table S9** Upregulated DEGs of *B. cinerea* in 12 h vs. control.

**Supplemental Table S10** Downregulated DEGs of *B. cinerea* in 12 h vs. control.

**Supplemental Table S11** DEGs of *B. cinerea* in pathogenesis in 12 h vs. control.

## Acknowledgements

This study was supported by the National Key Research and Development Program of China (No. 2019YFD1002004).

## Data availability statement

The data that support the findings of this study are available from the corresponding author upon reasonable request.

